# A molecular model of the mitochondrial genome segregation machinery in *Trypanosoma brucei*

**DOI:** 10.1101/188987

**Authors:** Anneliese Hoffmann, Sandro Käser, Martin Jakob, Simona Amodeo, Camille Peitsch, Jiří Týč, Sue Vaughan, Benoît Zuber, André Schneider, Torsten Ochsenreiter

## Abstract

In almost all eukaryotes mitochondria maintain their own genome. Despite the discovery more than 50 years ago still very little is known about how the genome is properly segregated during cell division. The protozoan parasite *Trypanosoma brucei* contains a single mitochondrion with a singular genome the kinetoplast DNA (kDNA). Electron microscopy studies revealed the tripartite attachment complex (TAC) to physically connect the kDNA to the basal body of the flagellum and to ensure proper segregation of the mitochondrial genome via the basal bodies movement, during cell cycle. Using super-resolution microscopy we precisely localize each of the currently known unique TAC components. We demonstrate that the TAC is assembled in a hierarchical order from the base of the flagellum towards the mitochondrial genome and that the assembly is not dependent on the kDNA itself. Based on biochemical analysis the TAC consists of several non-overlapping subcomplexes suggesting an overall size of the TAC exceeding 2.8 mDa. We furthermore demonstrate that the TAC has an impact on mitochondrial organelle positioning however is not required for proper organelle biogenesis or segregation.

**Significance Statement:** Mitochondrial genome replication and segregation are essential processes in most eukaryotic cells. While replication has been studied in some detail much less is known about the molecular machinery required distribute the replicated genomes. Using super-resolution microscopy in combination with molecular biology and biochemistry we show for the first time in which order the segregation machinery is assembled and that it is assembled *de novo* rather than in a semi conservative fashion in the single celled parasite *Trypanosoma brucei*. Furthermore, we demonstrate that the mitochondrial genome itself is not required for assembly to occur. It seems that the physical connection of the mitochondrial genome to cytoskeletal elements is a conserved feature in most eukaryotes, however the molecular components are highly diverse.

**Abbreviation:**

(EZF): Exclusion zone filaments
(ULF): Unilateral filament
(TAC): tripartite attachment complex
(OM): outer mitochondrial
(IM): inner mitochondrial
(BSF): bloodstream form
(PCF): procyclic form
(kDNA): kinetoplast DNA
(gRNA): guide RNA
(SBFSEM): Serial block face-scanning electron microscopy
(Tet): tetracyclin
(STED): Stimulated Emission Depletion

## Introduction

Mitochondria are key organelles in almost all organisms. Their ability to generate energy via oxidative phosphorylation depends on a small number of proteins that are encoded on the mitochondrial genome (1, 2). Thus, proper replication and segregation of the mitochondrial genome is essential for cell growth and healthy tissues. While many aspects of the replication have been studied in great detail the segregation of the organelle’s genome is less well understood. Trypanosomes are parasitic, single celled eukaryotes within the supergroup of the Excavates. They are the causative agent of number diseases including African sleeping sickness and Chagas disease in humans and Nagana in cattle. Their single large mitochondrion has attracted attention especially from structural biologists using electron microscopy. Associated with the organelle they discovered a large electron dense body the kinetoplast that turned out to contain DNA, which was eventually shown to be the organelles genome (3–6). In *Trypanosoma brucei* the mitochondrial DNA (kinetoplast DNA, kDNA) consists of 5000 plasmid like minicircles, diverse in sequence each encoding three to five guide RNAs (gRNAs) that are required to edit the cryptic transcripts from the 25 maxicircles, which are the homologous structures of mitochondrial genomes in many other well studied eukaryotes (7–9). Each minicircle is physically connected to three other minicircles and the maxicircles are interwoven in this network such that the overall structure of isolated kDNA resembles a knight’s chain mail (10). In *T. brucei* the genome is tightly packed into a disc-like structure of about 450x150 nm, localized in the kDNA pocket adjacent to the flagellum basal bodies (11). More than 30 proteins including several different polymerases and helicases are involved in the replication of the kDNA (10, 12). Post replication the genome is segregated into the developing daughter cells through the movement of the basal bodies (13). In 2003 an electron microscopy study by Ogbadoyi and coworkers revealed a filamentous structure connecting the basal body and the mitochondrial DNA that was named tripartite attachment complex (TAC) (14). The three parts of the structure are the exclusion zone filaments (EZF), named for the lack of cytoplasmic ribosomes, that ranges from the base of the flagellum to the outer mitochondrial (OM) membrane; the differentiated OM and inner mitochondrial (IM) membranes, which in this region are resistant to detergent treatment and the unilateral filaments (ULF) that connect the IM membrane to the kDNA (14). The ULF can be subdivided into a region with DNA enriched in basic proteins and a region without DNA with mostly acidic proteins (15).

A number of individual TAC components have been identified and characterized in recent years (Table 1). Starting with the most proximal component to the kDNA TAC102. TAC102 is a 102 kDa structural, basic (pI 9.2) protein with a mitochondrial import sequence in the C-terminal region, which is part of the ULF (16). Depletion of the protein initially leads to missegregation of the mitochondrial genome such that the daughter cell with the old basal body retains large parts of the kDNA while the daughter cell with the new basal body can only bind very small portions of the mitochondrial genome (16). Eventually, this leads to few cells with giant kinetoplasts and loss of the kDNA in the majority of the population. The function of TAC102 is restricted to mitochondrial genome segregation since loss of the protein has no impact on mitochondrial genome replication, organelle morphology, biogenesis or segregation (16). Consequently TAC102 is dispensable in a trypanosome cell line (γL262P) that is able to survive without mitochondrial genome, similar to the petite mutants in yeast (16–18). Further details of the molecular functions of TAC102 remain elusive. The first component of the TAC to be described, is p166 a 166 kDa large acidic (pI 5.1) protein, with a N-terminal mitochondrial targeting sequence that localizes to the region between the kDNA and the inner mitochondrial membrane (19). It contains a potential transmembrane domain that is not required for localization, however it remains unclear if it is required for proper function of the protein (19). Similar to TAC102, p166 is stably associated with the TAC in flagella isolated from cells using detergent and high salt conditions (16, 19). Interestingly, the p166 RNAi phenotype is quasi identical to the TAC102 RNAi phenotype. TAC40 is an OM membrane beta-barrel protein of the porin family similar to MDM10 from yeast (20). While the yeast MDM10 is involved in a number of different functions including the endoplasmatic reticulum mitochondrial encounter structure (ERMES) complex, nucleoid segregation and protein import machinery assembly (21–23), the function of TAC40 is restricted to mitochondrial genome segregation (20). Based on localization and biochemical purifications TAC40 is closely associated with TAC60 another OM protein with exclusive function in kDNA segregation. TAC65, another TAC component, which is not an integral mitochondrial OM protein biochemically interacts with pATOM36 (see below) and thus resides in the EZF (24). In the same region p197 was discovered during proteomics screens to characterize the basal body and bilobe structure of the flagellum (25). Similar to p166, p197 has been suggested to be a TAC component in procyclic form (PCF) parasites. For both proteins it remains unknown if they are also essential in bloodstream from (BSF) cells and if their function is restricted to mitochondrial genome segregation. Furthermore, Mab22 a monoclonal antibody against an unknown protein was identified to localize to the EZF and to the mature and pro-basal body (26). There are a number of additional proteins that either partially localize to the TAC region or to have an impact on proper mitochondrial genome segregation in trypanosomes. However, these proteins were also shown to be involved in functions other than genome segregation. This includes AEP1 a mitochondrial protein that results from alternative editing of mitochondrial COX3 transcripts and localizes to the TAC in isolated flagella (27). pATOM36, a OM protein with dual function in mitochondrial protein import machinery assembly and the kDNA segregation (24). Depletion of the mitochondrial acyl carrier protein (ACP), an enzyme of the fatty acid biosynthesis pathway leads to missegregated kinetoplasts in BSF but not PCF parasites, where a RNAi knockdown results in a cytochrome-mediated respiration defect (28, 29). The missegregation in BSF might be due to phospholipid composition changes in the mitochondrial membranes (28). As to why the two life cycle stages are affected differently remains unknown. Furthermore, TbCCD1 a member of the tubulinbinding cofactor C protein family that is involved in bilobe structure formation and proper connection of the TAC to the basal bodies (30). Finally, depletion of the Krebs cycle enzyme α-KDE2 results in an unequal distribution of the replicated kDNA suggesting a moonlight function for this well studied protein (31).

**Table 1:** TAC proteins and reagents

| Name | Gene ID | Reference | Information |
| --- | --- | --- | --- |
| BBA4 | – | (32) | - Decorates a protein of the basal- and pro-basal body<br>- Stably associates with isolated flagella |
|  |  | This study | - Loss of BBA4 after p197 RNAi |
| p166 | Tb927.11.3290 | (19) | - First discovered TAC component<br>- Transmembrane domain (residue 1440-1462)<br>- Localizes between kDNA and basal body (PCF)<br>- kDNA missegregation/growth retardation after RNAi (PCF)<br>- Stably associates with isolated flagella |
| | | This study | - Phenotype in BSF and not essential in BSF γL262P cells |
| Mab22 | – | (26) | - Decorates a protein in the EZF and mature/pro-basal body |
| p197 | Tb927.10.15750 | (25) | - Localizes between kDNA and basal body (PCF)<br>- kDNA missegregation/growth arrest after RNAi (PCF) |
| | | This study | - Localizes to the EZF<br>- Phenotype in BSF and not essential in BSF γL262P cells |
| TAC40 | Tb927.4.1610 | (20) | - Beta-barrel OM protein<br>- kDNA missegregation/growth arrest after RNAi (BSF, PCF)<br>- kDNA missegregation (BSF γL262P)<br>- Localizes between kDNA and basal body<br>- Stably associates with isolated flagella |
| TAC102 | Tb927.7.2390 | (16) | - Localizes to the ULF<br>- kDNA missegregation/growth arrest after RNAi (BSF, PCF)<br>- kDNA missegregation (BSF γL262P)<br>- Stably associates with isolated flagella |
| TAC65 | Tb927.5.830 | (24) | - Localizes to the EZF<br>- kDNA missegregation/growth arrest after RNAi (PCF, BSF)<br>- kDNA missegregation (BSF γL262P)<br>- Stably associates with isolated flagella |
|  |  | This study | - Localizes to the EZF |
| TAC60 | Tb927.7.1400 | (33) | - OM membrane protein with two transmembrane domains<br>- kDNA missegregation/growth arrest after RNAi (BSF, PCF)<br>- kDNA missegregation (BSF γL262P)<br>- Localizes between kDNA and basal body<br>- In complex with TAC40<br>- Stably associates with isolated flagella |

## Results

### p166 and p197 are essential TAC components in BSF cells

In order to verify that p166 and p197 are indeed essential components of the TAC in both major life cycle stages and to test if they are involved in other mitochondrial functions than genome maintenance we generated cell lines that allowed RNAi depletion in wild type BSF cells and a BSF cell line that is able to survive without mitochondrial genome but under wild type conditions retains the kDNA (γL262P, (17)).

RNAi targeting p166 in BSF (NYsm) parasites results in kDNA missegregation, eventually leading to a population of cells without kDNA. Growth of the population is affected three days post induction of RNAi (Fig. S 1A). However, no growth defect is detected in the γL262P cell line while maintaining the kDNA loss phenotype, suggesting the protein is only required in the context of mitochondrial genome segregation (Fig. S 1B). Similarly, RNAi targeting p197 leads to kDNA missegregation and growth defect in NYsm cells after two days of RNAi induction, while no growth defect is visible in the “petite mutant” of the BSF cells (Fig. S 2A B).

To investigate if the loss of p197 has an impact on basal body morphology we used transmission electron microscopy (TEM). Thin sections of the basal body did not show any obvious differences between wild type and p197 depleted cells indicating that the function of p197 is related to TAC biogenesis rather than basal body biogenesis (Fig. S 2C). In this context we also discovered that the protein or proteins decorated by the BBA4 antibody (recognizing an unknown protein lining the basal body (32)) disappear in the NYsm cell line upon p197 depletion, indicating that the protein/proteins are not required for proper growth and or basal body biogenesis and that BBA4 is potentially a TAC component (Fig. S 2D).

Thus p166 and p197 are both core components of the TAC with exclusive functions in kDNA maintenance.

### Relative order of the TAC proteins

To determine the relative order of the proteins within the TAC we measured the distance between the individual components using a combination of confocal and super-resolution microscopy (STED). The BSF cells were fixed and proteins of the TAC and the basal body were visualized using different antibodies as well as the kDNA was imaged using DAPI (Fig. 1A). YL1/2 an antibody targeting tyrosinated alpha tubulin and RP2, is located to the distal end of the basal body at the transitional fibers (Fig. 1A; (34–36)). The individual TAC components were either tagged with different protein tags (myc, PTP or HA) or in the case of TAC102 we used a monoclonal α-TAC102 antibody (16). The distance measurements between the TAC components and the kDNA were normalized to the distance of the kDNA to YL1/2 (see materials and methods; Fig. 1B). Based on our measurements, TAC102 is the kDNA most proximal currently known TAC component with a median relative distance of 0.268. It is followed by p166 with a relative distance of 0.352 to the kDNA. Next are TAC40 and TAC60 both of which biochemically localize in the OM membrane (20, 33) and accordingly both show a very similar median relative kDNA distance value of 0.423 (TAC40) and 0.425 (TAC60). The two remaining proteins TAC65 and p197 are both part of the EZF (24, 25) and show a relative distance of 0.440 and 0.481, respectively. Thus, p197 is the kDNA most distal currently characterized TAC protein. Only the unidentified protein recognized by the BBA4 antibody is further away from the kDNA than p197 with a median value of 0.519.

**Fig. 1:**
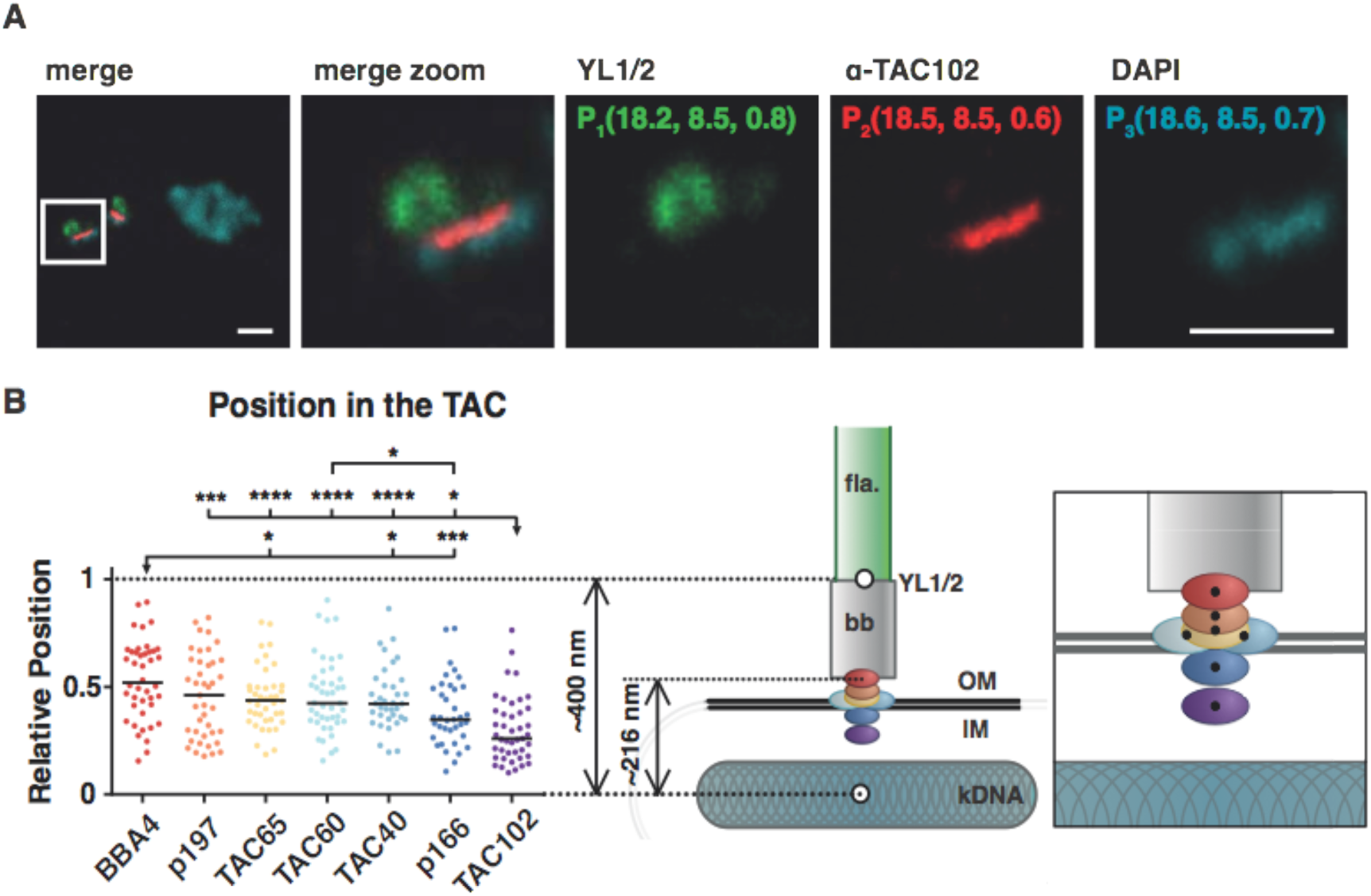
Relative position of the TAC components within the complex. A) A representative image of a 2K1N cell stained with DAPI for the nuclear and mitochondrial DNA (cyan) and YL1/2 for the basal body (green) is shown. Both channels were acquired with a confocal microscope. TAC102 is stained with a monoclonal antibody (red) and was acquired using STED microscopy. To measure the distance to the kDNA the center of mass was calculated by using the 3D object counter from Fiji. The xyz-coordinate per signal is shown as an example in the single color images. By using the Pythagorean theorem the distance could be reckoned. B) All measuring points for the relative position of the different components (BBA4 red, p197 orange, TAC65 yellow, TAC60 light blue, TAC40 blue, p166 dark blue, TAC102 purple) are indicated by dots and a black line indicates the median (36≤ N≤ 44). The model shows the relative position within the complex (right model). The flagellum (fla) is highlighted in green, the basal body (bb) in grey, the kDNA in cyan-grey and two black lines depict the mitochondrial membrane (OM, IM). A zoom in of the TAC components within the complex is shown, next to it. Scale bar 1 *μ*m.

### TAC assembles in a hierarchical order from the basal body

Based on the distance measurements we were able to test what impact the relative position of each component has on the overall assembly of the TAC structure. For this we applied RNAi against each of the TAC components and then analyzed after tetracyclin (Tet) induction the presence of remaining TAC proteins by epifluorescence microscopy. Depleting the kDNA most proximal TAC component, TAC102, leads to missegregation of kDNA and eventually loss of the mitochondrial genome in the majority of the cells, as described previously (16). However, despite the loss of kDNA no substantial changes in localization of the OM membrane protein TAC40 was detected (Fig. 2A). Thus, in the absence of TAC102 the localization of the more kDNA distal TAC component TAC40 remains unchanged. On the other hand, if we deplete p197 the kDNA most distal core component, we detect a similar kDNA loss phenotype as described for TAC102 and we also loose the epifluorescence signal for TAC40 (Fig. 2A). In order to verify the loss of TAC40 upon p197 RNAi we also probed for the protein in whole cell extracts by western blotting and found the overall levels of TAC40 to be decreased only marginally after 48 h of p197 depletion (Fig. 2B). This indicates that the loss of epifluorescence signal is likely due to misslocalization rather than degradation of tagged TAC40. In order to test all combinations of RNAi knockdown against a TAC component while analyzing the presence of the remaining TAC signals, we created 23 different cell lines. The results for immunofluorescence analysis are summarized in Fig. 2C and Fig. S 3 – Fig. S 8. Depletion of any currently known TAC component leads to the loss of proper localization of the downstream, more kDNA proximal TAC components at the epifluorescence microscopy level. Additionally, the two mitochondrial OM membrane proteins TAC40 and TAC60 also impact each other’s localization such that TAC40 RNAi leads to loss of TAC60 and vice versa. Furthermore TAC65, which is very close to TAC40 and TAC60 but is not outer mitochondrial membrane component (24) is also impacted by the depletion of TAC40 and TAC60.

**Fig. 2:**
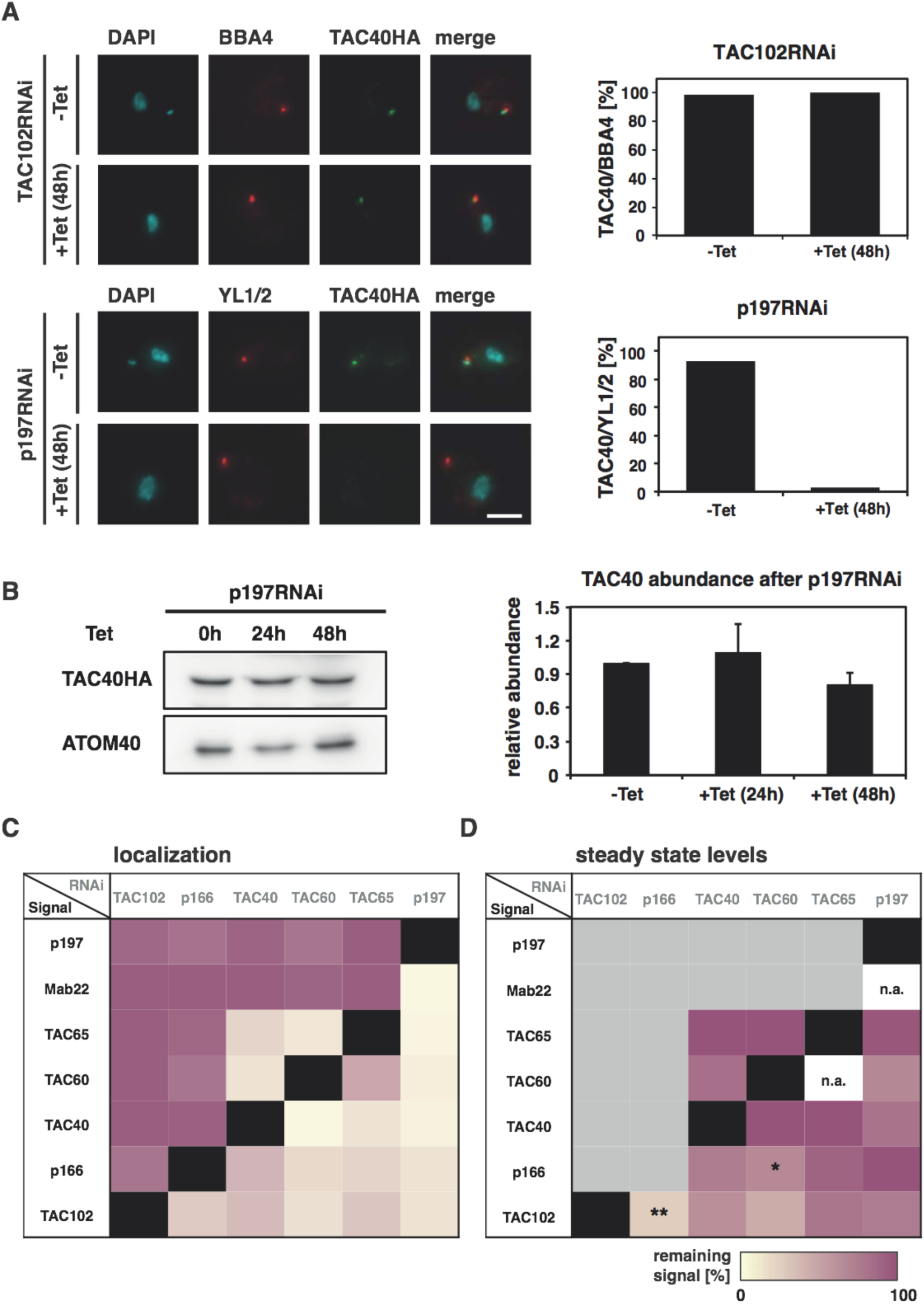
Depletion of a TAC protein and investigation of the remaining TAC components by Immunofluorescence and Western Blot analysis. A) Fluorescence microscopy images of TAC102 RNAi or p197 RNAi uninduced or 48 h Tet induced cells, stained with DAPI (DNA, cyan), BBA4 or YL1/2 (basal body, red) and TAC40HA (green). Next to it the quantitative analysis (N≈100). B) Protein abundance of TAC40 determined by western blotting from cells uninduced and 24 h, 48 h post p197 RNAi induction. Quantification of the protein abundance (N=3). ATOM40 abundance was used as a loading control. C) Quantification for each protein as described in A). Purple indicates that in 100% of the cells the signal can proper localize, whereas 0% is indicated by yellow. A black box shows that a knockdown of this component results in a loss of the signal for the same protein. D). Quantification for each protein as described in B). The percentage of the remaining signal after inducing for 48 h in comparison to the uninduced cells was calculated. Yellow indicates loss of signal in western blotting, purple indicates no changes. A black box shows that a knockdown of this component results in a loss of the signal for the same protein and a grey implements no investigation on western blot, since no changes occurred in localization; n/a not applicable. Calculating the p-value by two-tailed heteroscedastic t-test performed significance measurements (* p≤0.05; ** p≤0.01; *** p≤0.001; **** p≤0.0001). Scale bar 3 *μ*m.

When we evaluated the protein abundance level of the TAC components in the same cell lines by western blotting, we found that in general, loss of the epifluorescence signal did not correlate with loss of the protein by western blotting (Fig. 2D, Fig. S 9 – Fig. S 13). However, there are two exceptions. Depletion of p166 leads to a significant loss of the TAC102 protein (0.001<p<0.01) and depletion of TAC60 in the OM membrane leads to a significant loss of the p166 protein (0.01<p<0.05), which we suspect to reside at the IM membrane (19).

From this data we conclude that the TAC is built in a hierarchical order from the basal body towards the kDNA such that the kDNA proximal components require the upstream proteins for proper localization.

An alternative explanation of the data observed could be that the upstream components are not required for assembly but rather the lack of connection between the upstream and downstream components would lead to the observed phenotype. In this case we could expect that the downstream components accumulate on the old TAC structure since they cannot be distributed to the new daughter cell (Fig. 3A). We tested this model by depleting TAC40 and measured if TAC102 would accumulate at the old TAC structure that is associated with the enlarged kDNA as shown previously (Fig. 3B; (16)). Integrated density measurements showed no changes in TAC102 intensity 48 h post TAC40 depletion, thus supporting the hierarchical model of the TAC assembly (Fig. 3C).

**Fig. 3:**
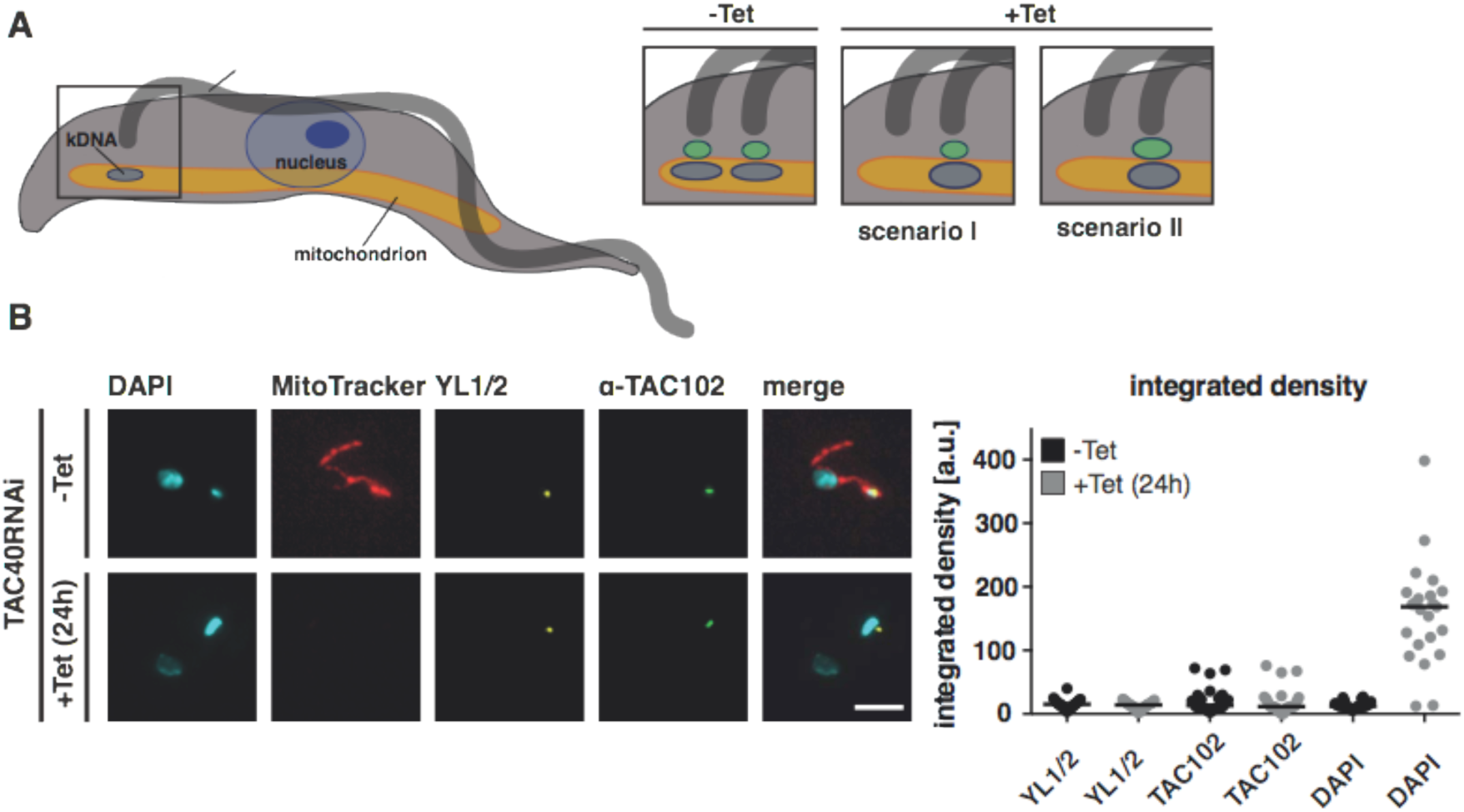
Downstream TAC component does not accumulate upon RNAi depletion. A) Model of a trypanosome with a zoom in of the TAC in uninduced and induceced 2K cells. Induced 2K cells show either no signal and no kDNA or an enlarged kDNA, which either is connected to a “normal” sized TAC (scenario I) or an enlarged one (scenario II). B) Uninduced (MitoTracker stained, red) and 24 h induced TAC40 RNAi cells were mixed and stained with YL1/2 (basal body, yellow), TAC102 (green) and DAPI (DNA, cyan). C) The integrated density of 1K1N uninduced and induced (enlarged kDNA) cells for YL1/2, TAC102 and DAPI was measured with Fiji (N_unind._=30, N_ind._= 23). Scale bar 3 *μ*m

### TAC consists of subcomplexes

Since the loss of epifluorescence microscopy detection was due to mislocalization rather than protein degradation, we wondered if the individual proteins might be assembled in subcomplexes that would prevent their proteolysis in the case of improper localization. In order to investigate this we applied blue native gel electrophoresis in combination with western blotting to characterize the TAC in BSF trypanosomes. Previously, TAC65 was shown to migrate in a 300 kDa complex in PCF (24). We could confirm the complex size of TAC65 in BSF cells. Under native conditions TAC102 migrates at around 440 kDa, while p166 is in a distinct complex larger than 670 kDa. The largest currently known complexes seem to be formed by TAC40 (between 500 and 900 kDa) and TAC60 that forms several distinct bands two of which migrate larger than 670 kDa (Fig. 4A). Based on the extraction and native running conditions there are at least five different subcomplexes that can be identified in BSF cells. There is very little overlap in complex size of the different TAC components. When we deplete TAC40 and then probe for p166 on blue native gels we find the complex to be largely unchanged in apparent size and abundance (Fig. 4B), the same could be observed for TAC60 after depleting p197 (Fig. S 14). Thus confirming our hypothesis that depletion of a basal body proximal TAC component leads to a mislocalization of the downstream partners but the individual subcomplex remain unaffected.

**Fig. 4:**
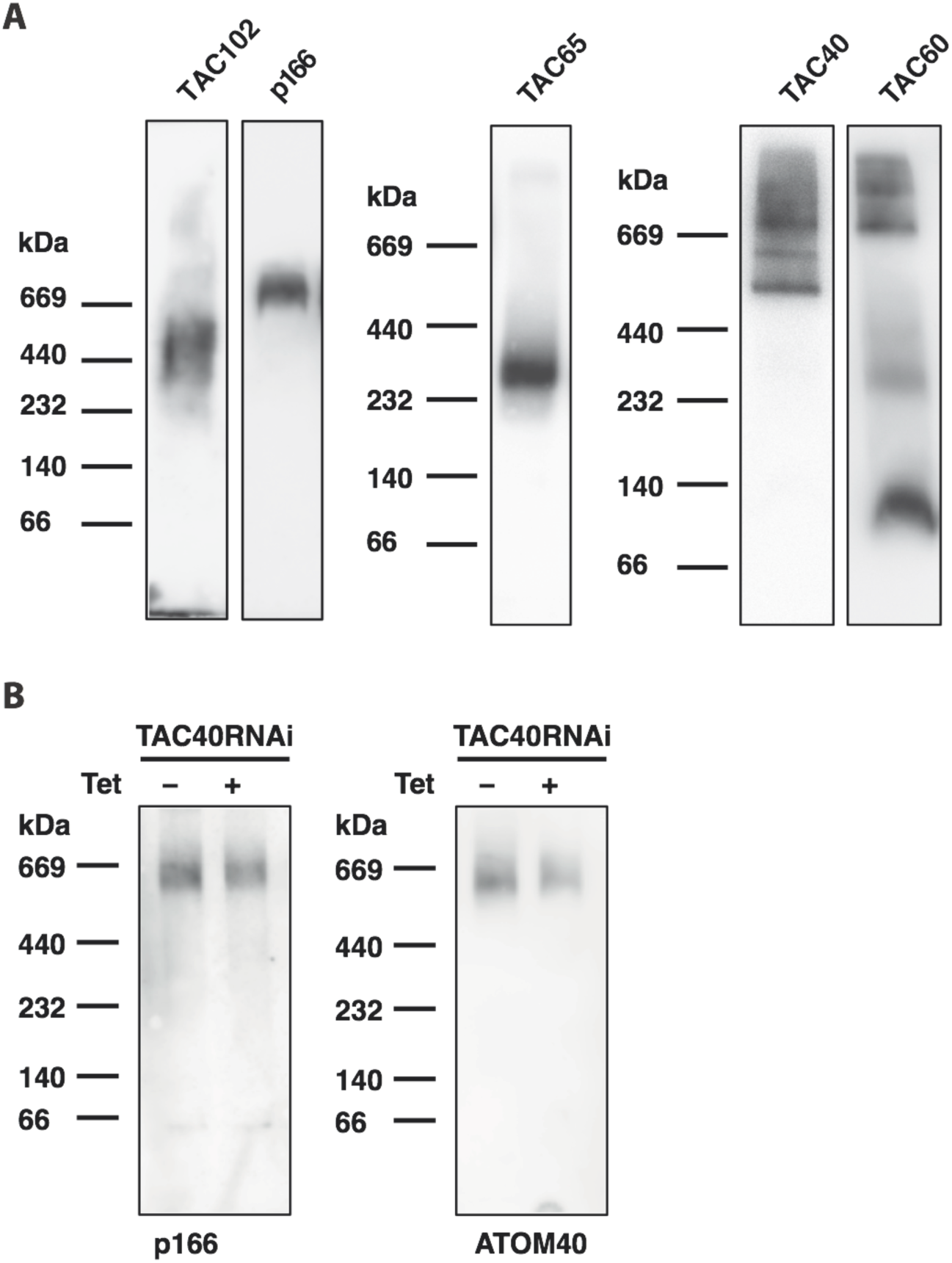
Complex formation of individual TAC components analyzed on blue native PAGE. A) Western blot from blue native PAGE of mitochondrial protein fraction of BSF cells decorated with the following antibodies: anti-TAC102, PAP for p166, myc for TAC60 and TAC65 or HA for TAC40. B) Same as A) from TAC40 RNAi induced cells (day 0 and day 2). The membrane was either decorated with an anti-p166 antibody or anti-ATOM40, as loading control.

### kDNA is not required for TAC assembly

The depletion of individual TAC proteins demonstrated that upstream components are required for proper localization of the downstream kDNA proximal elements. However, it remained unclear if the TAC itself can re-assemble in the absence of kDNA. To investigate this, we used the trypanosome cell line γL262P (17). In this cell line we depleted the basal body proximal TAC component p197 by RNAi for five days until ≥99% of the cells had lost their mitochondrial genome while maintaining wild type growth rates (Fig. 5A, B). We then probed for TAC components representing the three regions of the TAC, Mab22 (EZF), tagged TAC40 (OM membrane) and TAC102 (ULF) by epifluorescence microscopy (Fig. 5C). At day five post RNAi induction 14% of the cells showed a weak and 42% a mislocalized TAC102 signal, for the remaining 44% of the cells no signal was detected (Fig. 5D). Costaining with MitoTracker confirmed the misslocalized signal of TAC102 to be mitochondrial (Fig. 5E). At the same time post p197 depletion 94% of the cells showed no signal and 6% a mislocalised signal for TAC40 and Mab22 the EZF marker could not be detected in any cells (Fig. 5D). We then removed Tet from the medium to stop depletion of p197 and investigated the signal for TAC102, TAC40 and Mab22. Two days post recovery TAC102 localized to the correct position adjacent to the basal body marker (YL1/2) in 98% of the cells. For Mab22 and TAC40 99% (92% normal, 7% weak) and 74% (40% normal, 34% weak) of the cells showed a signal, respectively. We also depleted p166 in the γL262P cell line (Fig. S 15A, B). In this case, 44% of the cells had lost a proper signal for TAC102 (13% normal signal, 40% weak signal, 3% mislocalized signal) after 5 days of induction (Fig. S 15C, D). One day post recovery, 62% show a weak signal and after two days 100% of the cells had recovered the wild type TAC102 signal. In contrast upon p166 depletion no changes for the EZF marker Mab22 could be detected (Fig. S 15D). Thus the TAC can assemble de novo in the absence of the mitochondrial genome.

**Fig. 5:**
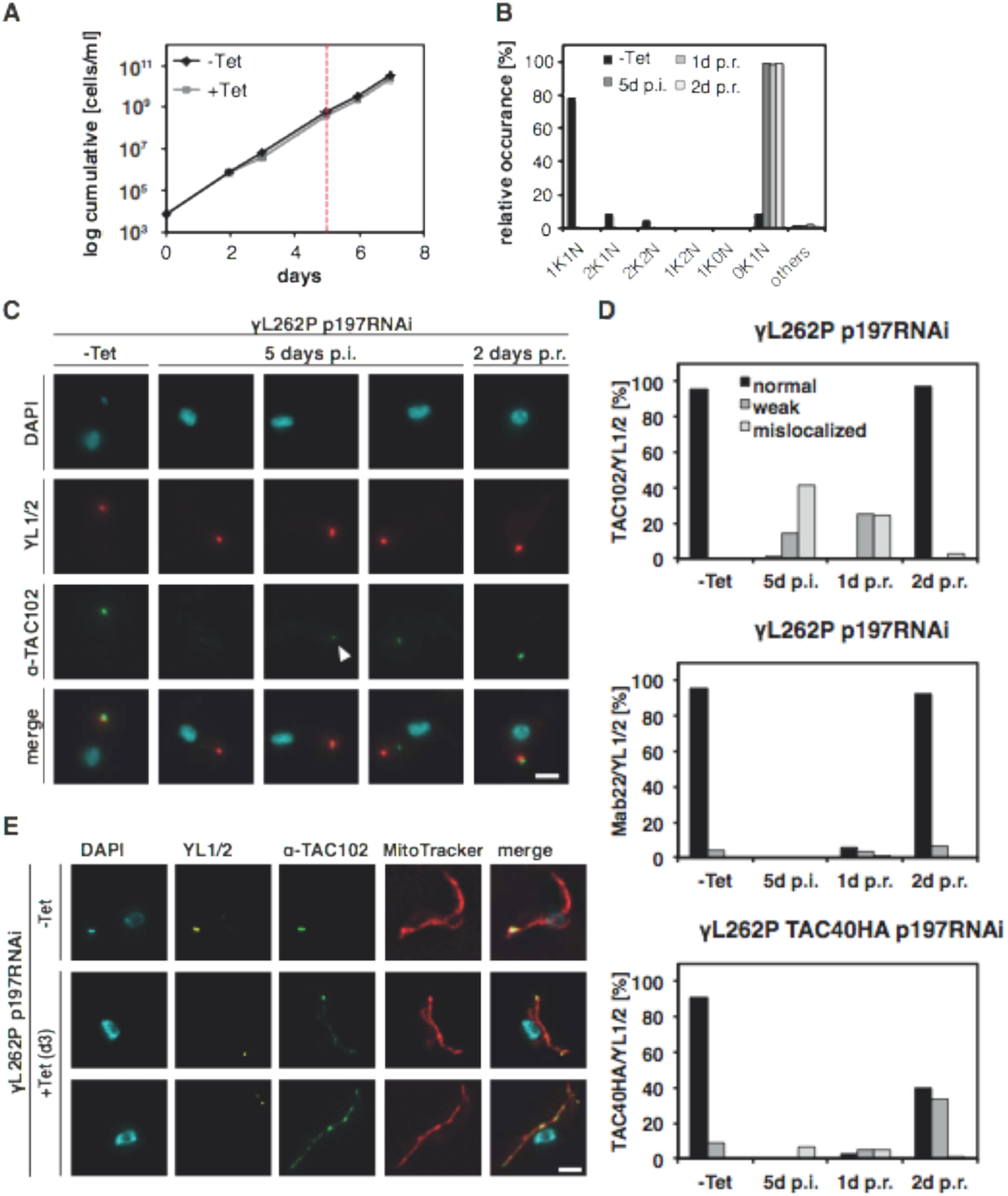
Recovery of the TAC after releasing p197RNAi in γL262P. cells A) Growth curve after 197 RNAi induction in γL262P cells with Tet. After 5 days of induction cells were washed and cultivated in medium without additional Tet (red line). B) The percentage of the relative occurrence of kDNA and nucleus in p197 RNAi induced and uninduced cells (N≈100). C) TAC102 (green), YL1/2 (basal body, red) and DAPI (DNA, cyan) stained cells after 0 and 5 days of Tet induction (post induction p.i.) and two days post recovery (p.r.) are shown. White arrowhead points to the weak signal. D) Quantitative analysis of TAC102 (N≥106), Mab22 (N≥93) or TAC40HA (N≥119) in uninduced (-Tet), 5 days induced (5d p.i.) and 1 or 2 days after removing Tet (1d p.r., 2d p.r.) in γL262P p197 RNAi cells. A black bar indicates a correct localized signal; a grey bar indicates a weak signal and a light grey a mislocalized signal. E) p197 RNAi γL262P cells were Tet induced for 3 days and stained with MitoTracker (Mitochondria, red), TAC102 (green) and YL1/2 (basal body, yellow). The DNA was stained with DAPI (cyan). Scale bar 2 *μ*m.

### Timing of TAC assembly during the cell cycle

The next question we wanted to address was the timing of TAC assembly during the cell cycle. Previous studies have timed mitochondrial genome is replicated after basal body duplication, but prior to the nuclear genome (11, 37, 38). During the replication, the unit size kDNA grows into a bilobed structure that is subsequently segregated into two kinetoplasts. One can easily detect four different stages of kDNA replication: (i) the unit size kDNA, (ii) the enlarged kDNA, (iii) the bilobed kDNA and the (iv) segregated kDNA connected via the nabelschnur (38). We followed individual markers of the three TAC regions, relative to the mitochondrial genome replication and segregation (Fig. 6A, Fig. S 16). Prior to kDNA replication the vast majority of cells show one signal for each of the three markers corresponding to one TAC structure being present. During the replication of the kDNA first the basal body proximal components like BBA4 and later TAC40 are assembled into a new TAC, clearly separated from the old structure. The last component to be assembled based on our data is TAC102 the kDNA most proximal TAC component (Fig. 6B, C).

**Fig. 6:**
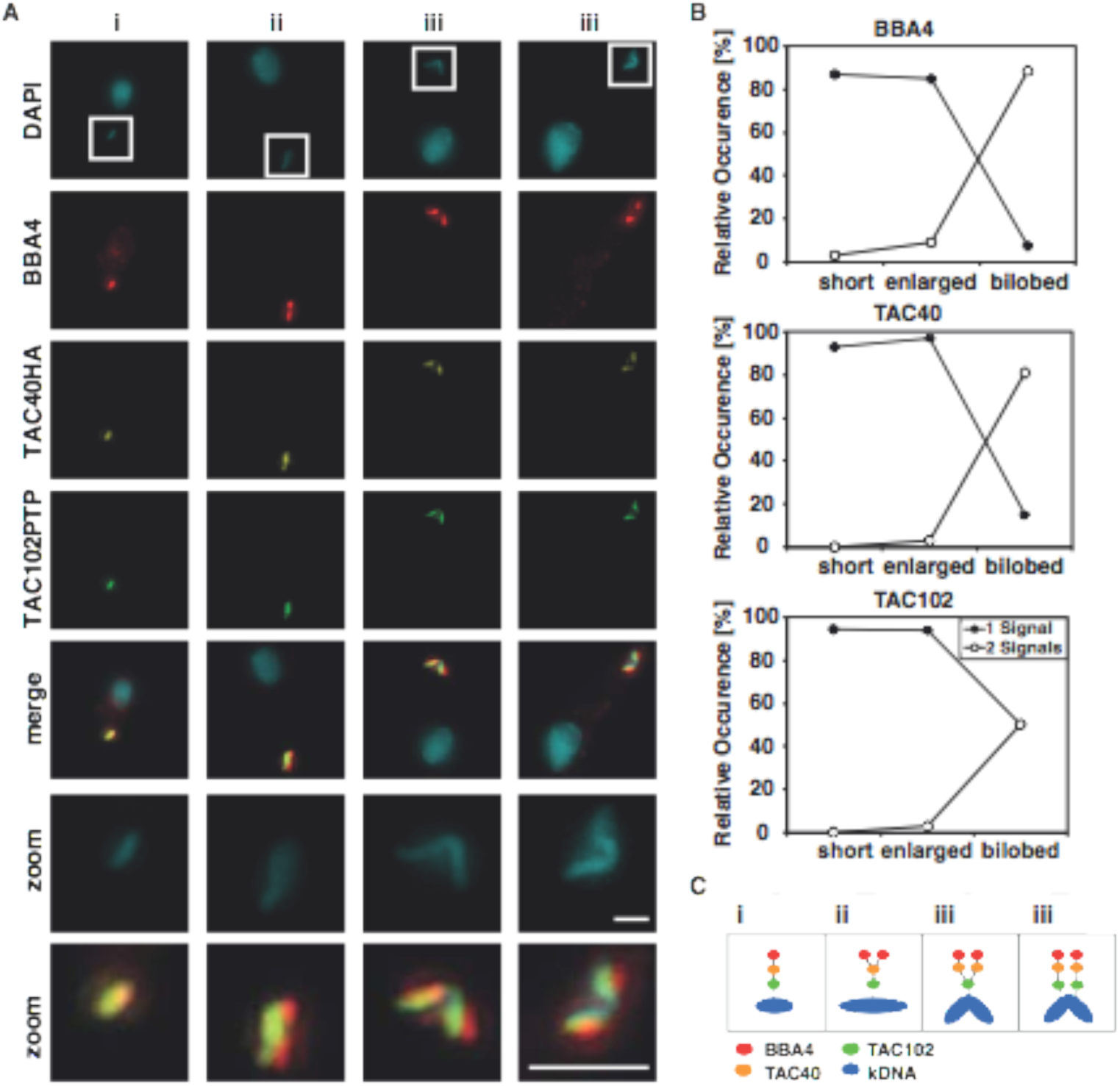
Dynamics of the TAC proteins during the kDNA replication. A) Tagged BSF cells were stained with DAPI (DNA, cyan), BBA4 (red), anti-HA for TAC40HA (yellow) and anti-ProteinA for TAC102PTP (green) to investigate the replication of the TAC in more detail. B) Cells were analyzed, divided into stage i (short), ii (enlarged) and iii (bilobed) (38) and the number of signals per staining were counted (N=125). C) An schematically model illustrates the TAC replication. Scale bar 1 *μ*m

### Physical connection of the basal body to the mitochondrial membranes

Based on our current hierarchy model we would predict that severing the TAC in the EZF would lead to a change in localization of the mitochondrion relative to the basal body, while severing the connection in the ULF should have no effect on the basal body-mitochondrial positioning. In order to test this model we depleted either p197 or TAC102 and used serial block face-scanning electron microscopy (SBFSEM) to analyze the distance between the basal body and the mitochondrial outer membrane, compared to the wild type situation (Fig. 7A, B). After TAC102 depletion no changes in the distance of the basal body to the mitochondrial membrane could be observed, while the median distance after p197 knockdown significantly increases from 124 nm to 283 nm (Fig. 7C). Thus, indeed the TAC complex also holds the posterior region of the mitochondrion in place.

**Fig. 7:**
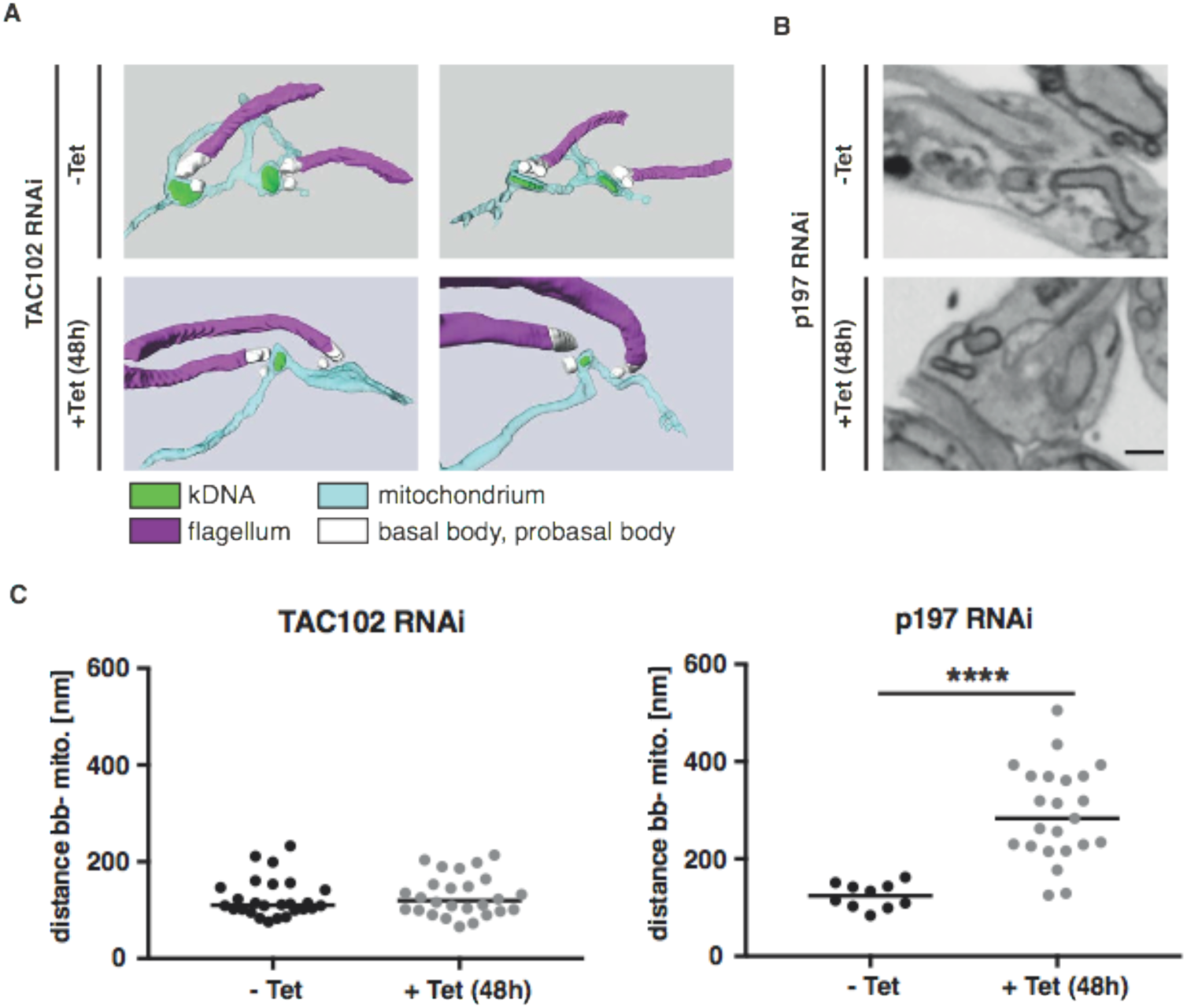
Distance formation between the mitochondria and the basal body after TAC102 RNAi or p197 RNAi by using Serial block face-scanning electron microscopy (SBFSEM) A) 3D reconstructions of TAC102 RNAi uninduced and 48 h induced cells are shown. B) 48 h induced and uniduced p197 RNAi electron microscopy images are shown. C) The distance between the basal body and the mitochondrial membrane was measured for TAC102 (N= 26) and p197 (N_unind._=10; N_ind._=23) uninduced and 48 h induced cells. Calculating the p-value by two-tailed heteroscedastic t-test performed significance measurements (* p≤0.05; ** p≤0.01; *** p≤0.001; **** p≤0.0001).

## Discussion

In this study we describe the architecture and assembly of the mitochondrial genome segregation machinery also named the tripartite attachment complex (TAC) in trypanosomes. The TAC is a large structure about twice the diameter of a nuclear pore complex (> 200 nm) and encompasses three regions in the cell, the cytoplasm, the outer and inner mitochondrial membranes and the mitochondrial matrix (14, 39). It provides the physical connection between the base of the flagellum and the kDNA that allow the segregation of the mitochondrial genomes in concert with the movement of the basal bodies during the cell cycle. Thus, similar to the centrioles that nucleate the microtubule organizing centers, which in turn are responsible to build the spindles that separate sister chromatids during mitosis, the basal bodies in *T. brucei* organize the mitochondrial segregation machinery (40, 41)

Based on super-resolution microscopy we are able to localize each component within a particular region of the TAC (Fig. 1). One exception is Mab22, a monoclonal antibody that only recognizes its antigen if the cells are strongly extracted with detergent. Since this procedure is potentially changing the structure within the cells we did not include this antibody in the distance measurements. Overall the distance measurements are in good agreement with the available biochemical data for TAC40 TAC60, and TAC65. TAC40 and TAC60 are both mitochondrial OM membrane proteins and thus should be positioned between p166 a protein with a canonical mitochondrial targeting sequence and p197, which is clearly non-mitochondrial (19, 20, 25, 33).TAC65 is not a mitochondrial protein and thus should localize between the OM proteins TAC40/TAC60 and p197 (24). Furthermore, the positioning of TAC102 (pI 9.2) in a region close to the kDNA is consistent with the electron microscopy data that demonstrated a region of basic proteins in the ULF close to the kDNA (15). Similarly, our super-resolution data places p166 (pI 5.1) further away from the kDNA than TAC102 in a region that has been reported to contain acidic proteins close to the inner mitochondrial membrane (15).

In order to test if TAC biogenesis is organized in a particular hierarchy or if assembly occurs independently in different regions of the complex we depleted each TAC component and subsequently checked if and how assembly of the TAC and its components is affected (Fig. 2). The results from these experiments are not consistent with an assembly starting at the kDNA since depletion of the kDNA most proximal component TAC102 has no impact on the localization of any other TAC component that is at greater distance to the kDNA. In the random assembly model we would expect the signal for a kDNA proximal component to accumulate if a distal component is depleted. However, as we demonstrated for TAC102 after TAC40 depletion this is not the case; while the TAC102 signal remains at the old basal body kDNA connection it does not significantly increase in intensity (Fig. 3). Thus, only the hierarchical model where assembly of the TAC starts at the base of the flagellum and extends through the two-mitochondrial membranes to the kDNA, is consistent with our current data.

This model is also supported by the imaging data that describe the TAC biogenesis during the replication cycle of the kDNA (Fig. 6). Clearly the TAC components close to the basal body appear first in two separate structures while it happens only just prior to segregation for TAC102 the kDNA most proximal protein. Furthermore, the recovery experiments in the petite yeast-like trypanosome cell line showing TAC assembly in the absence of kDNA also supports the hierarchical model since even lack of the mitochondrial genome does not impact TAC assembly (Fig. 5). Interestingly, the complex involved in mitochondrial genome maintenance and segregation in yeast that has been named two membrane spanning structure (TMS) can also assemble in the absence of the mitochondrial genome (42). The TMS was identified through co-localization of the mitochondrial matrix protein Mgm101 with a subset of the mitochondrial nucleoids and the outer membrane protein Mmm1 by immunofluorescence microscopy (42) and while it seems clear that both proteins together are required for proper mitochondrial DNA maintenance the actual connection to the cytoskeleton remains unknown.

Including previous work and this study more than ten proteins have been described to be involved in TAC biogenesis in BSF and PCF trypanosomes. Six of these proteins, p197 and TAC65 in the EZF, TAC40 and TAC60 in the OM membrane, p166 and TAC102 in the ULF seem to exclusively function in genome maintenance (16, 19, 20, 24, 25, 33).The strategy to exclusively employ a number of different proteins for mitochondrial genome segregation seems unique to trypanosomes. Other model systems rely on proteins with multiple functions. In yeast MDM10/MDM12 and Mgm101 for example are involved in mitochondrial genome segregation but also in multiple processes like ER mitochondrial connections (MDM10/12), oxidative mtDNA damage repair (Mgm101) and protein import (MDM10) (2, 43, 44). A similar situation can be observed in mammalian cells where the two proteins Mfn1/2 are involved in mitochondrial ER junction formation as well as nucleoid maintenance (45). As to why trypanosomes have developed such an elaborate system of specialized proteins remains unknown but we can speculate that the single unit nature of the kDNA and its complexity and size were important factors in this development. For TAC102 we could show that the protein is present in all currently sequenced Kinetoplastea (16), however a detailed analysis has not been done for the remaining TAC components.

Native gel electrophoresis identified high molecular weight complexes associated with the proteins TAC102, p166, TAC65 as well as TAC40 and TAC60 in BSF cells (Fig. 4). For TAC65 a complex of similar size had previously been shown in the PCF of the parasite supporting that the TAC is conserved between the two life cycle stages (24). However, there is very little overlap between the different complexes indicating that even under the mild detergent conditions several subcomplexes are isolated. The largest subcomplexes seem to be in the mitochondrial outer membrane where TAC40 and TAC60 have been shown to interact in biochemical pull down experiments. The two proteins of the ULF p166 and TAC102 do not seem to reside in the same subcomplex, which is also supported by the significant distance between the two proteins that place the very basic TAC102 in the region close to the kDNA, while the acidic p166 is potentially IM membrane associated. Gluenz and colleagues, previously demonstrated the presence of two biochemically different regions within the ULF (15).

Based on this information TAC102 is in close proximity to the kDNA, however we have no evidence for direct interactions between TAC102 and the kDNA. While we could clearly demonstrate that the EZF TAC components are required for proper positioning of the kDNA pocket close to the base of the flagellum (Fig. 7), the lack of proper positioning does not seem to influence proper organelle division and thus a separate mechanism for the distribution of the organelle during cell division must exist. Arguably the TAC is now the best described mitochondrial genome segregation machinery, however there are many aspects that still remain elusive including the nature of the filamentous structure in the EZF, the connection between inner and outer mitochondrial membrane as well as the connection to the mitochondrial genome itself just to name a few.

## Materials and Methods

### Cell culture

All experiments were done either with the BSF *T. brucei* strain New York single marker (NYsm) (46) or the γL262P cell line (17). Cells were grown in HMI-9 medium supplemented with 10% FCS (47) at 37°C and 5% CO_2_. Depending on the cell line 2.5 *μ*g/ml geneticin 0.5 *μ*g/ml puromycin, 2.5 *μ*g/ml phleomycin, 5 *μ*g/ml blasticidin or 2.5 *μ*g/ml hygromycin were added. RNAi was induced through the addition of 1 *μ*g/ml tetracyclin.

### Transfection with different plasmids

20 or 40 million cells were transfected with 8-10 *μ*g of plasmid in 90 mM Na_3_PO_4_, 5 mM KCl, 0.15 mM CaCl_2_, 50 mM HEPES, pH 7.3 transfection buffer by electroporation using Amaxa Nucleofector II program X-001 (48, 49). The p166 RNAi was targeted against the ORF (3452-3952 bp) of the Tb927.11.3290 gene and p197 RNAi against the ORF (2546-3083 bp) of Tb927.10.15750. The p166 RNAi construct was created by using the gateway cloning system (50). RNAi against p197 was generated by using the phleomycin containing pLew100-derived stemloop vectors (51). Using HindIII/XbaI and XhoI/BamHI as restriction enzymes with a stuffer of 460 bp formed the hairpin. Before transfection, both plasmids were linearized by NotI. Cells containing TAC40 RNAi, TAC60 RNAi or TAC65 RNAi were described previously (20, 24, 33).For tagging p197 and p166, the ORF region 4-1264 bp or the ORF region 3204-4503 bp was PCR-amplified from genomic DNA, respectively. The PCR product for p197 was digested with NotI and ApaI and ligated into these sites of pN-PURO-PTP (52). The resulting plasmid was linearized with BmgBI before transfection. The p166 PCR product was digested with ApaI and EagI and ligated into ApaI and NotI of pC-PTP-PURO (52). This vector is a derivative of pC-PTP-NEO in which the neomycin resistance gene was replaced by the ORF of the puromycin resistance gene via NdeI and BstBI. The final plasmid was linearized with FspAI and transfected. Cells containing TAC40 HA, TAC65 myc or TAC60 myc tag were described previously (20, 24, 33). All used primers are summarized in Table S 1.

### Immunofluorescence microscopy

Cells were spread on a slide and fixed with 4% PFA in PBS (137 mM NaCl, 2.7 mM KCl, 10 mM Na_2_HPO_4_, 2 mM KH_2_PO_4_, pH 7.4) for 4 min. After washing with PBS the cells were permeabilized with 0.2% TritonX-100 for 5 min. After 30 min of blocking with blocking solution (4% BSA in PBS), slides were incubated for 45 or 60 min with the primary antibody followed from the secondary antibody incubation for 45 or 60 min at room temperature. The antibodies were diluted in blocking solution. For a double staining either the primary antibodies and then the secondary antibodies were mixed together, or they were used one after the other with an additional blocking between the first secondary and second primary antibody. All used antibodies are summarized in Table S 2. For the rat α-TAC102 antibody, the secondary antibody had to be diluted 1:500. Cells were mounted with ProLong™ Gold Antifade Mountant with DAPI (Molecular Probes). Acquisition was performed with the epifluorescence DM5500 microscope from Leica or a DFC360 FX monochrome camera (Leica Microsystrems) mounted on a DMI6000B microscope (Leica Microsystems). Image analysis was done using LASX software (Leica Microsystems), ImageJ and Imaris.

### MitoTracker staining

Cells were stained in medium with 200 nM MitoTracker™ Red CMXRos (Molecular Probes) as described in the manufactures instructions. For mixing unstained and stained cells additional washing steps are crucial. A subsequent immunofluorescence staining was performed as described above. All images were taken from the same microscopic slide using the same acquisition settings. Images were deconvolved using the deconvolution software from Leica (LAS AF 2.6.1.7314). In ImageJ maximum intensity z-projections were made. Binary masks were generated manually for each channel separately using the same linear signal intensity threshold values. The integrated density (area * mean grey intensity) of each particle was evaluated using the “measure particles” functionality provided by ImageJ. In the settings, the original 16-bit signals were redirected to obtain the values.

### Dynamics of the TAC

Immunofluorescence analysis was performed as described above. Before blocking with 4%BSA in PBS, an additional incubation step for 30 min with Image-iT™ FX Signal Enhancer (Thermo Fisher) with a following PBS/T washing step was implemented. In ImageJ maximum intensity z-projections were made. Binary masks were generated manually for each channel separately using the same linear signal intensity threshold values. Kinetoplasts were recognized automatically as binary particles inside the DAPI channel. The center of mass was used to generate a squared cropping mask, big enough to contain the particles from all channels. The binary image stacks were quantified in a subsequent step. Based on the area size and binary shape descriptors, three classes of kDNA were assigned; short, enlarged and bilobed shaped. The number of particles were counted for each kDNA. The combination of the number of particles were outputted and automatically grouped and summarized for each kDNA class.

### Stimulated Emission Depletion (STED) microscopy and distance measurements

Cover glasses (#1.5) were glow discharged for 30 seconds with the FEMTO SCIENCE CUTE discharger. Cells were spread on the cover glass and the fixing, permeabilization and staining was performed as described above. Images were acquired by using the SP8 STED microscope (Leica) as z-stacks with a z-step size of 120 nm. In order to minimize differences that might occur during the cell cycle we only used cells that already had duplicated and segregate the mitochondrial genome (2K1N, 2K2N). To obtain the distance of the TAC components to the kDNA the centre of mass was determined by using the 3D object counter in Fiji. With this plug-in it is possible to reckon the xyz coordinates for the centre of mass of an object. The distance between two objects can be calculated by using the Pythagorean theorem. To achieve the relative position, the measurements of the TAC component to the kDNA distance were normalized to the distance of the kDNA to the basal body.

### Cytoskeleton extraction

For the antibodies Mab22 a cytoskeleton extraction needed to be performed. For this, total cells were washed with PBS and spread on a slide. After removing the liquid cells were incubated for 1 minute with extraction buffer (100 mM PIPES pH 6.8, 1 mM MgCl_2_) containing 0.05% Nonidet P-40. Afterwards the cells were washed with extraction buffer and the staining was completed as described above.

### Western Blot analysis

5x10^6^ cells were mixed with 1x Laemmli buffer (0.4% SDS, 12 mM Tris-HCl pH 6.8, 4.8% glycerol, 1% β-mercaptoethanol, bromophenol blue in PBS) and loaded per lane on a 6%, 8% or 10% SDS-polyacrylamide gel. Blocking (5% or 10% milk in PBS/T) after transferring onto a PVDF membrane was performed for 1 h at room temperature. Primary antibodies were incubated for 1 h at room temperature or over night at 4°C, besides PAP which was incubated for 30 min at room temperature. Secondary antibodies were incubated for 1 h at room temperature. All used antibodies are summarized in Table S 2 and were diluted in blocking solution. Acquisition was performed with the ODYSSEY Infrared imaging system (LI-COR), the LAS1000 (Fuji Medical Systems) or the Amersham Imager 600 (GE Helathcare).

### Blue native gel electrophoresis

For an enriched mitochondria fraction 0.025% or 0.015% digitonin in SoTE (0.6 M Sorbitol, 20 mM Tris-HCl pH7.5, 2 mM EDTA) was used. With this fraction it was proceeded as described previously (53). Instead of 1.5% digitonin 1% was used and incubated for 15 minutes on ice. After a centrifugation step the supernatant was loaded on a native gradient gel. Afterwards the gel was soaked in SDS-buffer (25 mM Tris, 192 mM glycine, 0.1% SDS) and transferred onto a PVDF membrane by semi-dry western blotting. The decoration with the antibodies was performed as described above.

### Transmission electron microscopy of thin sections

After harvesting, the cells were treated as described previously (16). Thin sections were scanned with a transmission electron microscope (FEI Morgani, Tungsten cathode). The microscope was equipped with a digital camera (Morada, 12 megapixel, Soft Imaging System) and the AnalySIS iTEM image analysis software.

### Serial block face-scanning electron microscopy

Sample preparation, data processing and analysis was performed as described previously for TAC102 RNAi uninduced and 48 h induced cells (24). Serial images of the block face were recorded at an accelerating voltage of 4 kV, a spot size of 1 and pressure of 0.33 (+Tet) or 0.3 (-Tet) Torr. Pixel size and the dwell time for each micrograph was 5 nm and 3.2 *μ*s respectively and slice thickness was 100 nm. For p197 RNAi 48 h induced and uninduced cells, block staining, dehydration and embedding were performed as described previously (11). The shortest distance between the basal body and the mitochondrial membrane was measured manually with IMOD. To do so, the block was orientated to get the basal body and the flagellum as well as the kDNA pocket in one plane. This was not possible for the induced samples since the kDNA pocket was not preserved. In order to measure the basal body and the flagellum were oriented in one plane and the nearest mitochondrial tube was used for measuring. Because of this, the measurement had to be done throughout several slices.

## Acknowledgment

We would like to thank Keith Gull for the BBA4 and YL1/2 antibodies as well as Derrick Robinson for Mab22. For technical assistance we acknowledge Bernd Schimanski, Adolfo Odriozola, Evelyne Vonwyl and Nicolas Niklaus. Imaging was performed with devices supported by the Microscopy Imaging Center (MIC) of the University of Bern, Switzerland and the Bioimaging Unit at Oxford Brookes University, UK. For financial support we thank the Novartis Foundation and the Berne University Research Foundation. Research in the lab of André Schneider was supported by grant 138355 and in part by the NCCR “RNA & Disease” both funded by the Swiss National Science Foundation. In the lab of Benoît Zuber, the grant 163761 by the Swiss National Science Foundation supported the research.

